## Supplementary Figures for "FBXO24 modulates mRNA alternative splicing and MIWI degradation and is required for normal sperm formation and piRNA production"

Zhiming Li, et al.

### Supplementary Figures

A

| | Human | Mouse | Rat | Chimpanzee | Cat | Bovine | Horse | Dog | Xenopus |
|---|---|---|---|---|---|---|---|---|---|
| Human | | 71.5% | 71.4% | 78.7% | 72.5% | 72.1% | 72.5% | 72.9% | 37.6% |
| Mouse | 71.5% | | 78.1% | 71.5% | 75.0% | 73.9% | 74.0% | 74.6% | 38.5% |
| Rat | 71.4% | 78.1% | | 73.9% | 75.4% | 76.5% | 74.9% | 75.1% | 38.8% |
| Chimpanzee | 78.7% | 71.5% | 73.9% | | 73.9% | 74.7% | 73.0% | 73.3% | 38.1% |
| Cat | 72.5% | 75.0% | 75.4% | 73.9% | | 76.5% | 77.8% | 79.2% | 38.8% |
| Bovine | 72.1% | 73.9% | 76.5% | 74.7% | 76.5% | | 76.1% | 76.8% | 39.5% |
| Horse | 72.5% | 74.0% | 74.9% | 73.0% | 77.8% | 76.1% | | 78.3% | 38.6% |
| Dog | 72.9% | 74.6% | 75.1% | 73.3% | 79.2% | 76.8% | 78.3% | | 39.2% |
| Xenopus | 37.6% | 38.5% | 38.8% | 38.1% | 38.8% | 39.5% | 38.6% | 39.2% | |

B

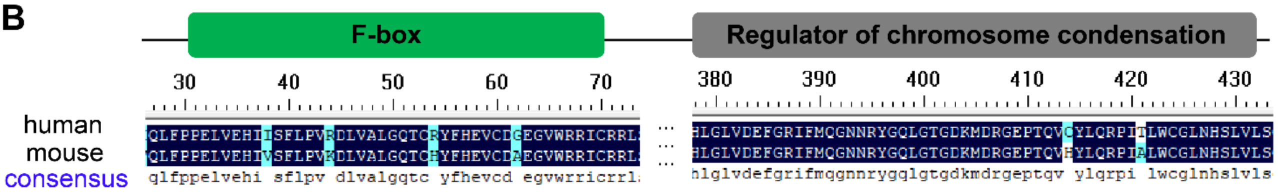


C

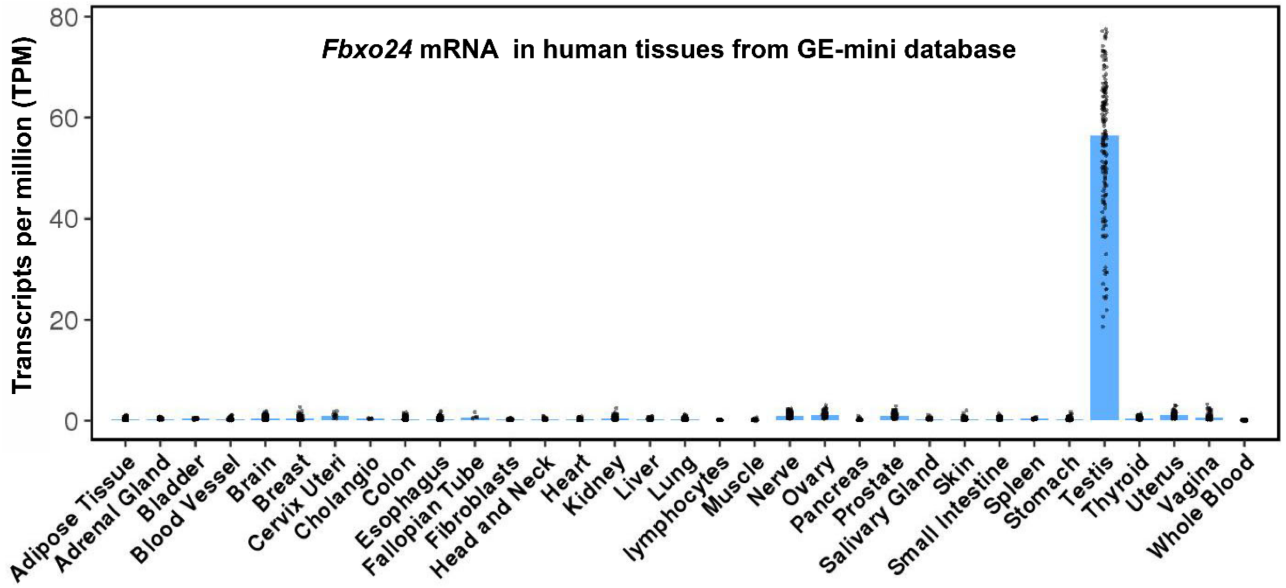


**Figure S1. Expression profiles of FBXO24 in mice and human.** (A) A high degree of conservation of FBXO24 in amino acid sequences among 9 species. (B) Amino acid sequence similarity of F-box and regulator of chromatin condensation (RCC1) of FBXO24 protein in mouse and human. (C) mRNA levels of FBXO24 in multiple human organs from GE-mini database.

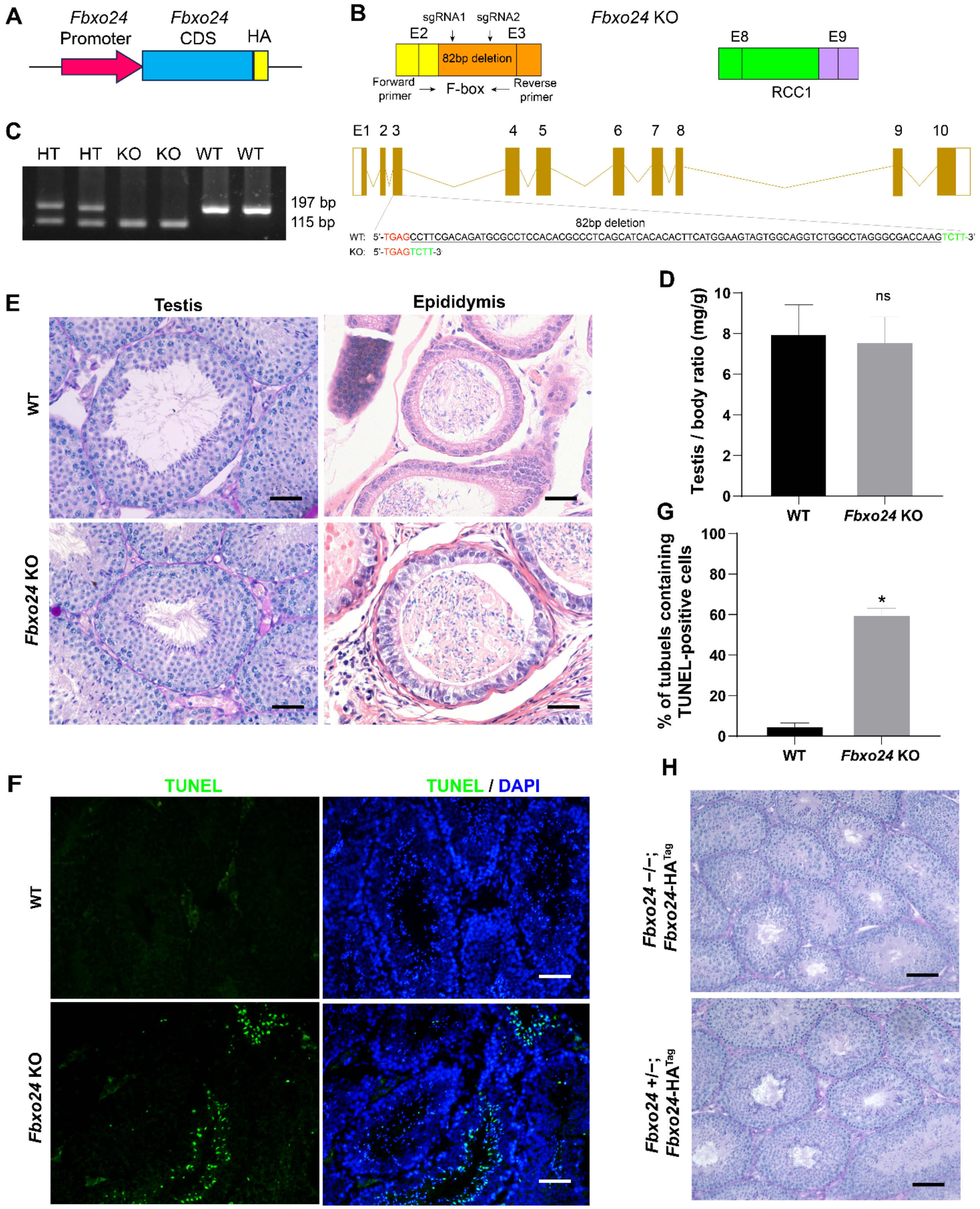


**Figure S2. Spermiogenesis was defective in FBXO24-deficient mice.** (A) Schematic representation of transgenic cassette. (B) Diagram illustrating the CRISPR/Cas9 targeting strategy, including position and sequence of guide RNAs (sgRNAs). E, exon. RCC1, Regulator of chromosome condensation. (C) Examples of PCR genotyping of the FBXO24 mutated region in WT (wild type), HT (heterozygote) and KO (knockout) mice. (D) Testis/body-weight ratio of WT and *Fbxo24* KO (n = 3/group) mice at 8-week-old. Data are mean ± S.D. ns, not significant. (E) Representative histological section images of testis and epididymis obtained from *Fbxo24* KO mice and WT mice stained with PAS and H&E, respectively. Scale bars, 50 mm. (F) TUNEL analysis of WT and *Fbxo24* KO testis are shown. Apoptotic cells were labeled by TUNEL staining (green). Scale bar, 50 µm. (G) Comparison of TUNEL-positive seminiferous tubules in WT and *Fbxo24* KO testis (n = 3/group). Error bars represent mean ± S.D. *, $p < 0.05$. (H) PAS staining for testicular sections of 8-week-old *Fbxo24*−/−; *Fbxo24*-$HA^{Tag}$ mice. Scale bar =50 µm.

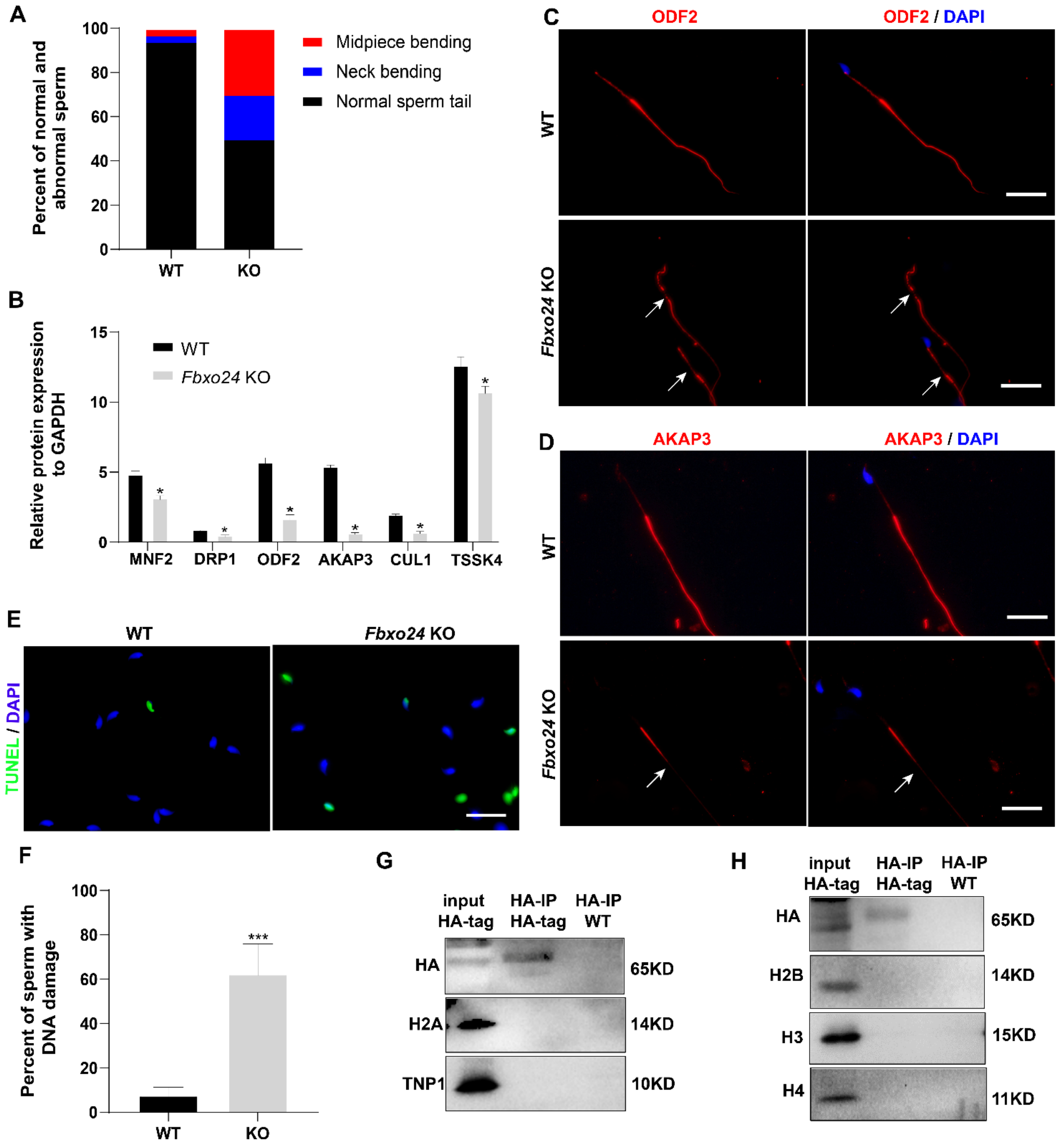


**Figure S3. Sperm morphology analysis in FBXO24-deficient mice.** (A) Percentage of morphologically normal and abnormal spermatozoa in WT and *Fbxo24* KO sperm. (B) Quantification of protein levels of MFN2, DPR1, ODF2, AKAP3, CUL1, and TSSK4 in WT and *Fbxo24* KO sperm. Error bars represent mean ± S.D., n = 3. *, $p < 0.05$. (C) ODF2 and (D) AKAP3 immunofluorescent signals in the flagellum of WT and *Fbxo24* KO sperm. The arrows indicated the week or absent staining of the immunofluorescent signal. Scale bar, 50 μm. (E) DNA damage was analyzed by TUNEL assay in WT and *Fbxo24* KO sperm. Scale bar, 50 μm. (F) Percentage of apoptotic cells in WT and *Fbxo24* KO sperm. Error bars represent mean ± S.D., n = 3. ***, $p < 0.001$. (G) The testis lysate of Fbxo24-HA tagged mice were immunoprecipitated with anti-HA beads. Western blots were used to detect the histones and transition protein expression.

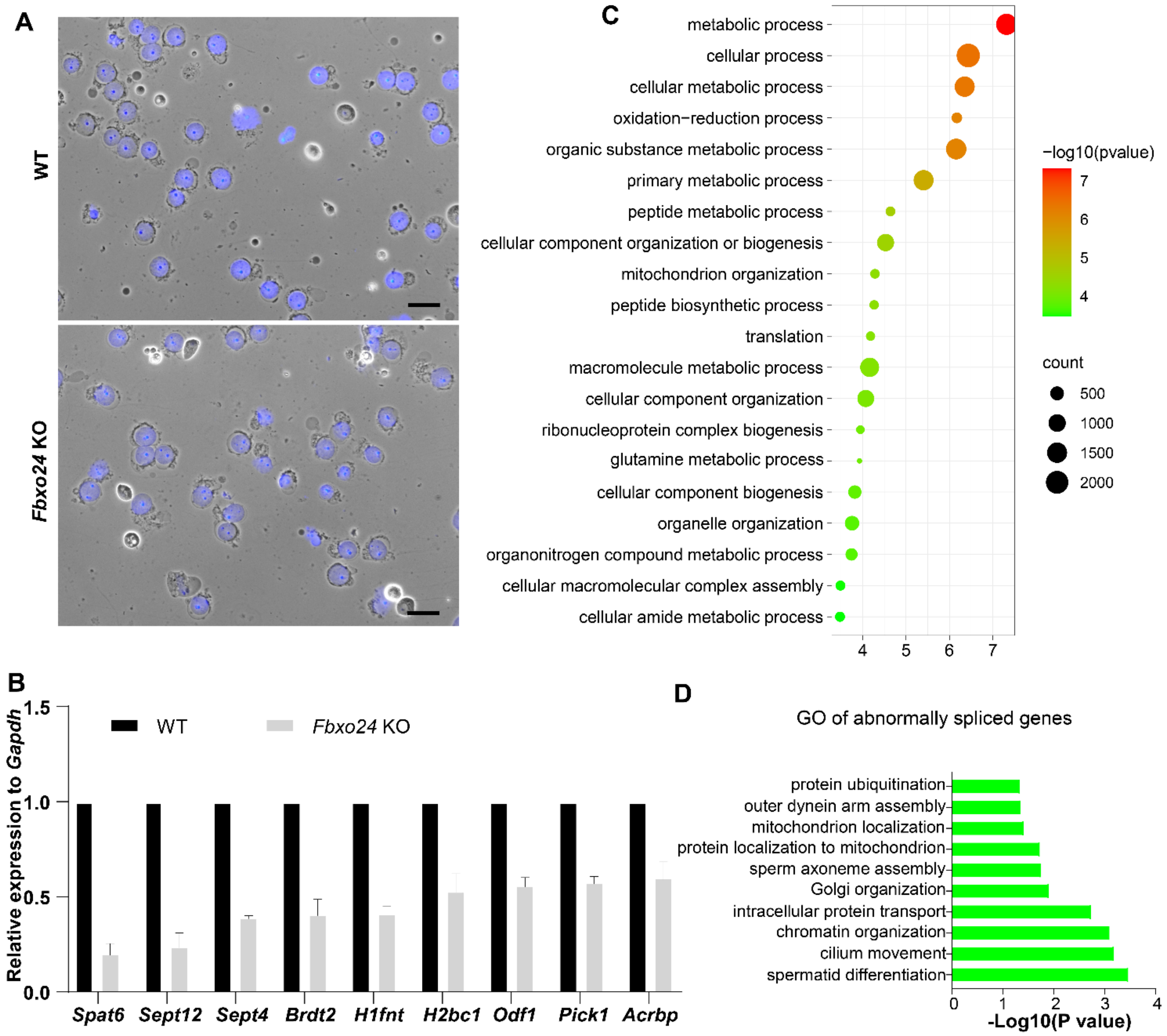


**Figure S4. Global gene expression altered in round spermatids of FBXO24-deficient mice.** (A) The purity and morphology of isolated round spermatids can be determined from bright-field images merged with DAPI nuclear staining (blue). Scale bar =50 µm. (B) qPCR validation of some genes in the testis of WT and *Fbxo24* KO adult mice. The data are represented as mean ± S.D. *, p< 0.05. n = 3. (C) Gene ontology of the top 20 GO terms of the up-regulated genes in *Fbxo24* KO round spermatids. (D) Gene ontology of the abnormally spliced genes in *Fbxo24* KO round spermatids.

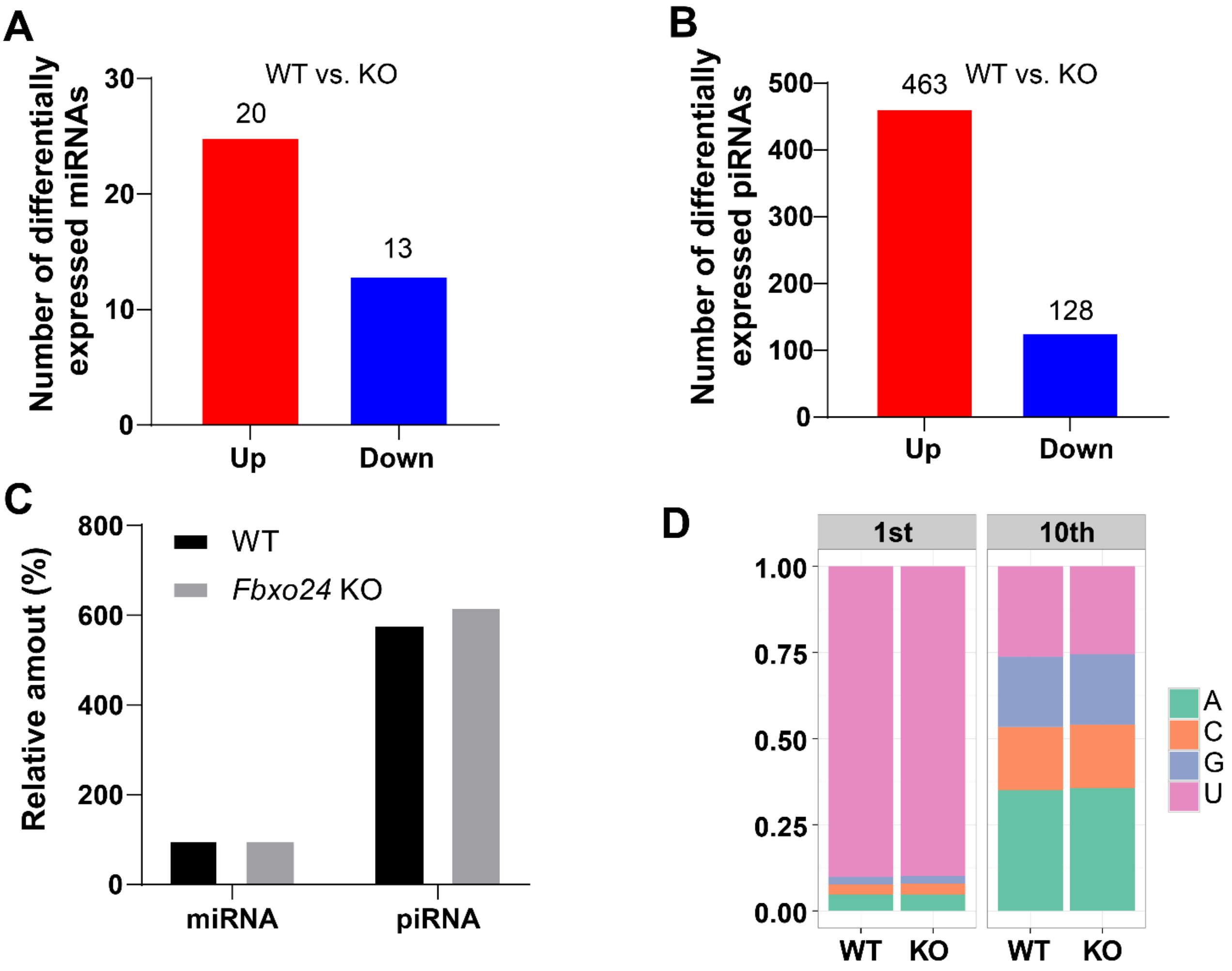


**Figure S5. miRNA and piRNA expression analysis in the FBXO24-deficient mice.** (A) Graph bars showing the number of differentially expressed miRNA in the testis of *Fbxo24* KO mice with 8-week-old (n =3 / group). Significantly regulated genes have a *p*-value of < 0.05 and fold change of > 2. The number of up- and down-regulated miRNA are indicated. (B) Graph bars showing the number of differentially expressed piRNA in the testis of *Fbxo24* KO mice with 8-week-old (n =3 / group). The number of up- and down-regulated piRNA are indicated. (C) Ratios of total piRNA in KO vs. WT after the normalization with miRNA counts. (D) Ratios of the 1st and 10th nucleotides of the repeat-associated piRNA.
